# Field validation of an eDNA assay for the endangered white-clawed crayfish *Austropotamobius pallipes*

**DOI:** 10.1101/562710

**Authors:** Siobhán Atkinson, Jeanette E.L. Carlsson, Bernard Ball, Mary Kelly-Quinn, Jens Carlsson

## Abstract

The white-clawed crayfish *Austropotamobius pallipes* has undergone extensive declines within its native range in the last century. Because of its threatened status, European legislation requires the species to be regularly monitored and that Special Areas of Conservation (SACs) be designated for it. Knowledge on the distribution of this species is vital for addressing these needs. This study presents an environmental (e)DNA assay to detect *A. pallipes* in water samples, based on the mitochondrial cytochrome oxidase I (COI) gene, utilizing species-specific primers, a minor groove binding (MGB) probe and quantitative PCR. The results of this study indicate that eDNA is an effective tool for detecting *A. pallipes* in a lotic system, and could provide a valuable, non-invasive method for determining the distribution of this species.

## Introduction

The white-clawed crayfish *Austropotamobius pallipes* is a relatively large, long-lived (>10 years) crustacean that inhabits both rivers and lakes (Reynolds et al. 2010). They require alkaline conditions for survival and are commonly found in waterbodies overlying carboniferous limestone bedrock (Lucey and McGarrigle 1987). *A. pallipes* is one of the five indigenous crayfish species in Europe (Holdich et al. 2009). This once abundant species has, however, become greatly reduced or locally extinct across large parts of its native range during the last century (Grandjean et al. 1997). Pollution (Demers and Reynolds 2002, Lyons and Kelly-Quinn 2003), habitat loss and disease (Matthews and Reynolds 1992) have contribution to this decline. Of particular concern is the crayfish plague, caused by the fungus *Aphanomyces astaci* (Holdich et al. 2009). *A. pallipes* possess no resistance to this fungus, so eradication of entire populations of *A. pallipes* is possible following an outbreak (Reynolds et al. 2010). Despite this, Ireland is considered one of the few remaining strongholds for *A. pallipes* within Europe. One of the reasons for this is that the invasive signal crayfish *Pacifastacus leniusculus*, which can impose wide-ranging impacts on ecosystems and act as a vector for the crayfish plague (Vaeßen and Hollert 2015), has not been reported in Ireland to date. However, the crayfish plague has reached Ireland’s rivers, and large changes in the distribution of *A. pallipes* have been attributed, at least in part, to past plague outbreaks (Demers et al. 2005). Additionally, there is at present an outbreak of the crayfish plague in Ireland (National Biodiversity Data Centre 2018), so more than ever there is a need for non-invasive sampling methods that do not have the potential to further spread the crayfish plague.

Because of its vital role in freshwater ecosystem functioning and its threatened status (Matthews and Reynolds 1992), *A. pallipes* is afforded protection under the EU Habitats Directive and is listed under Annexes II and V. This means Ireland is required to regularly monitor *A. pallipes*, and to designate Special Areas of Conservation (SACs) for the species under Natura 2000 (Reynolds et al. 2010). Knowledge on the geographic distribution of *A. pallipes* is essential for implementing these conservation measures. Traditional survey methods include night viewing with a strong torch, modified quadrat samplers (DiStefano et al. 2003), kick-sampling, baited traps and enclosures (Byrne and Lynch 1999) and SCUBA diving (Matthews and Reynolds 1992). While many of these methods are effective, they are time-consuming, potentially costly, and may not be effective when *A. pallipes* occur in low abundance. Furthermore, as white-clawed crayfish are a protected species in Ireland, several surveying methodologies require a licence from the National Parks and Wildlife Service. Certainly, a low-cost, non-invasive and effective methodology for assessing the distribution of *A. pallipes* is warranted. A promising new tool that has been shown to be both effective at detecting species and to save significant time in the field (Sigsgaard et al. 2015) is environmental DNA (eDNA) analysis. Environmental DNA is the collective term for DNA present freely in the environment (Taberlet et al. 2012, Thomsen and Willerslev 2015). The analysis has been successful in detecting species that occur in low abundance such as rare or endangered species (Mächler et al. 2014, Laramie et al. 2015, Sigsgaard et al. 2015, Boothroyd et al. 2016, Carlsson et al. 2017), or invasive alien species (Jerde et al. 2011, Goldberg et al. 2013, Fujiwara et al. 2016). Futhermore, eDNA assays have been successfully deployed for other crayfish species including *Astacus astacus, Astacus leptodactylus, P. leniusculus* (Agersnap et al. 2017, Larson et al. 2017), *Orconectes rusticus* (Dougherty et al. 2016, Larson et al. 2017), *Procambarus clarkii* (Cai et al. 2017) and *Faxonius eupunctus* (Rice et al. 2018).

The aim of this study was to develop an MGB based qPCR assay to detect the presence of the white-clawed crayfish, *A. pallipes* in environmental samples, and to test the reliability of the assay by comparing the results with field observation data.

## Methods

### Study sites and crayfish distribution data

Eight sampling locations within seven different rivers were selected for field validation of the assay. This research was carried out as part of a wider study assessing the impact of river obstacles on Atlantic salmon *Salmo salar.* An obstacle, (weir, ford crossing or bridge apron), was therefore present in the river at each sampling location and eDNA samples were collected both above and below the structure. Out of the eight sites sampled, four were in rivers with SAC status in which *A. pallipes* featured as a species of interest. Study rivers within the River Barrow and River Nore SAC included the Delour and Dinin rivers and study rivers within the Lower River Suir SAC included the Multeen and Duag rivers. Although the Burren River is a tributary of the River Barrow, it does not have SAC status.

The presence of *A. pallipes* in these rivers was confirmed using data provided by the Irish Environmental Protection Agency (EPA). These data are point data based on casual field observations made of *A. pallipes* by EPA staff whilst undertaking routine biological monitoring between the years 2007 and 2016 (W. Trodd, Irish Environmental Protection Agency, personal communication). In addition, it was possible at some sites to confirm the presence of crayfish by field observations made while electrofishing for *S. salar* and *S. trutta.* For the purposes of this study, *A. pallipes* observations were extrapolated to the wider subcatchment level. This meant that even if only one field observation of *A. pallipes* occurred in a subcatchment, it was assumed that there were likely to be *A. pallipes* in the rest of that subcatchment. This was considered the most meaningful way to present the data, because the EPA were not specifically looking for *A. pallipes* when their observations were made, so there is no certainty that *A. pallipes* did not occur in other parts of the subcatchment.

### eDNA qPCR assay development

Species-specific primers (forward primer: 5’-GGG TTA GTG GAG AGA GGG GT -3’, and reverse primer 5’-AAT CCC CAG ATC CAC AGA CG -3’) and a 5’-6-FAM labelled TaqMan^®^ minor groove binding probe (5’-TCA GCT ATT GCC CAC GCA -3’) for *A. pallipes*, which targeted a locus within the mitochondrial cytochrome oxidase I (COI) region, were designed in Primer Express 3.0 (Applied Biosystems-Roche, Branchburg, NJ). Including primers, the total amplicon size was 96 base pairs. To verify the species specificity for the *A. pallipes* assay in silico, probe and primer sequences were queried against the National Centre for Biotechnology Information (NCBI – http://www.ncbi.nlm.nih.gov/) nucleotide database with BLASTn (Basic Local Alignment Search Tool). The assay was tested in vitro with freshwater species including brown trout *S. trutta*, sea lamprey *Petromyzon marinus*, Atlantic salmon *S. salar*, allis shad *Alsoa alosa* and twaite shad *Alosa fallax.* The qPCR assay was optimized using tissue extracted from *A. pallipes.*

### eDNA collection, filtering and extraction

Water samples for eDNA analysis were collected in sterilized 2 L PET bottles from two locations at each study site, above and below the river obstacle, and filtered on site using a peristaltic pump. To avoid contamination, the water samples were taken in an upstream direction, prior to any individuals entering the river. At each sampling location, three replicate water samples were collected, one from each of the left margin, right margin and the center of the river channel. In addition, one negative field control consisting of ddH20 was also filtered. This resulted in a total number of six water samples for eDNA analysis and two field controls collected per study site. All water samples were filtered through 47 mm glass microfiber filters (1.5 μm). Filters were stored in 2.0 mL Eppendorf tubes and subsequently frozen at -20°C. To reduce contamination risk, all eDNA work was performed in a dedicated Low Copy DNA laboratory. A modified CTAB (cetyltrimethylammonium bromide) protocol (Möller et al. 1992) was employed for eDNA extractions. Briefly, one-half of the filter was transferred to a 2.0 mL Eppendorf tube. A total volume of 750 μL CTAB buffer (100 mM Tris-HCL, 20 mM EDTA, 1.4 M NaCl, 2% CTAB), and 7 μL of Proteinase K (20 mg mL^−1^) were added to the tube. Samples were then vortexed for 10 seconds followed by incubation at 56°C for 2 hours, after which 750 μL of phenol/chloroform/isoamyl alcohol (25:25:1 v/v) was added. The contents of the tube were manually mixed for 15 seconds and subsequently centrifuged (11,000 x g, 20 min). A new tube contained 750 μL of chloroform/isoamyl alcohol (24:1 v/v) was prepared, and the aqueous phase was transferred to it. The manual mixing followed by centrifugation steps were repeated, and the aqueous phase was again transferred to a new tube. The DNA was precipitated using one volume of isopropanol alcohol that was added to the aqueous phase and incubated at -20°C for 1 hour followed by centrifugation (11,000 x g, 20 min). The resulting pellets were washed with 750 μL of 70% ethanol and centrifuged (11,000 x g, 5 min). The ethanol was removed, and care was taken to ensure that the pellet remained in the tube. The tubes containing pellets were dried in a heat block (50°C, 5 min) followed by resuspension in molecular-grade ddH2O.

### eDNA assay deployment

An Applied Biosystems ViiA™ 7 (Life Technologies, Inc., Applied Biosystems, Foster City, California, U.S.A.) quantitative thermocycler was used for eDNA amplification and quantification. Briefly, the PCR profile consisted of 50°C for 2 min and 95°C for 10 min, followed by 40 cycles between 95°C for 15 s and 60°C for 1 min. Standard curves were generated from DNA originating from *A. pallipes* tissue extracts (quantified with NanoDrop^®^- 1000, Thermo Scientific, Wilmington, DE). This served as both a positive control and as a means of calculating the concentration of *A. pallipes* eDNA in each sample. Seven 10:1 serial dilutions ranging from 2.9 ng μL^−1^ to 2.9 x 10^−6^ ng μL^−1^ were used for the standard curve. Each qPCR reaction was carried out in a total volume of 30 μL, consisting of 15 μL of TaqMan^®^ Environmental Master Mix 2.0 (Life Technologies, Applied Biosystems, Foster City, CA), 3 μL of each primer (final concentration of 2 μM), probe (final concentration of 2 μM), 3 μL eDNA/tissue extracted DNA template and 3 μL ddH2O. Individual standard curves were generated for each qPCR plate (y = -3.37x + 20.589, efficiency = 98.04 %, R^2^ = 0.996 (1), y = – 3.442x + 20.392, efficiency = 95.22 %, R^2^ = 0.999 (2) and y = -3.441x + 20.723, efficiency = 95.26 %, R^2^ = 0.999 (3)). Each plate had 2 no-template controls (NTCs) to identify any contamination that may have occurred during preparation. Standard curve, field and control samples were quantified in triplicate (3 technical replicates). A positive detection was defined as being when at least 2 out of the 3 technical replicates contained amplifiable DNA with Cq differences not exceeding 0.5, and when the samples were within the detection range of the standard curve. If 1 out of 3 technical replicates showed Cq differences exceeding 0.5, the replicate was excluded from the study. In cases when the Cq value of 2 out of 3 technical replicates differed by more than 0.5 Cq, that specific dilution series or field replicate was removed from further study. As *S. trutta* was present in all rivers, both upstream and downstream of the obstacles, this species was used as a positive field control to test for the presence of amplifiable DNA in all samples. The *S. trutta* assay from previously published work (Gustavson et al. 2015) was used for this analysis. Three replicates per location with 1 technical replicate were used.

## Results and Discussion

The present study was successful in detecting *A. pallipes* in silico, in vitro and in situ. The assay was highly sensitive and could detect *A. pallipes* eDNA concentrations as low as 0.002 ng L^−1^ at Cq 38.2 (average over 3 technical replicates, standard deviation 0.000059 ng L^−1^). No cross-species amplification occurred when the assay was tested using conventional PCR visualized on agarose. Furthermore, zero amplification occurred in the NTCs and field controls. While all field samples analysed yielded detectable DNA with the positive field control (S. *trutta* assay), this was not the case for the *A. pallipes* assay. *A. pallipes* was only detected in 5 out of 8 sites (Table 1, Fig. 1). Within the Burren, Duag and Multeen rivers, positive detections were observed in all field replicates above and below the river obstacles. At the Dinin (Cretty Yard) site, positive detection was observed in 2 out of 3 field replicates above the obstacle, and 3 out of 3 field replicates below the obstacle. At the Dinin (Castlecomer) site, positive detection of *A. pallipes* occurred in only 1 field replicate, which was located above the obstacle (Table 1). The detection of *A. pallipes* eDNA above the obstacle but not below it was a surprising result. Intuitively, one would expect that *A. pallipes* would have been detected below the obstacle also, as eDNA travels downstream in a river system. It is worth noting that it was at this site that the lowest concentration of *A. pallipes* eDNA was detected (0.002 ng L^−1^), and amplification of eDNA occurred in only 1 out of 6 site replicates. It is possible that eDNA concentrations in the other field samples were too low for a positive detection.

**Table 1.**
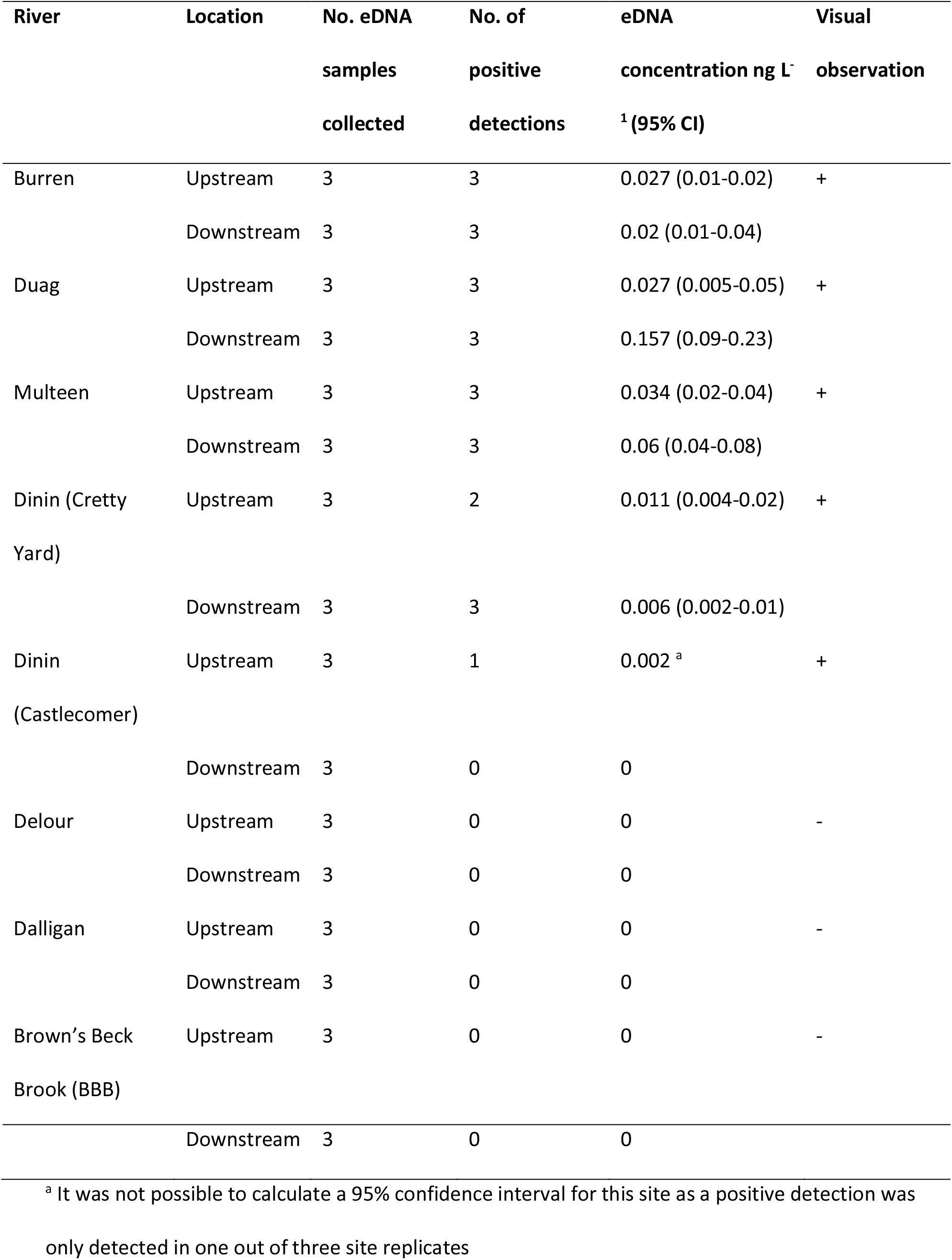
Environmental DNA concentrations (ng L^−1^) from the *A. pallipes* assay at each study river. Concentrations are based the average eDNA concentration in a minimum of 2 technical replicates and between 1 and 3 field replicates. Confidence intervals (95%) are given, where possible, for each location (upstream or downstream of the river obstacle).

**Figure 1.**
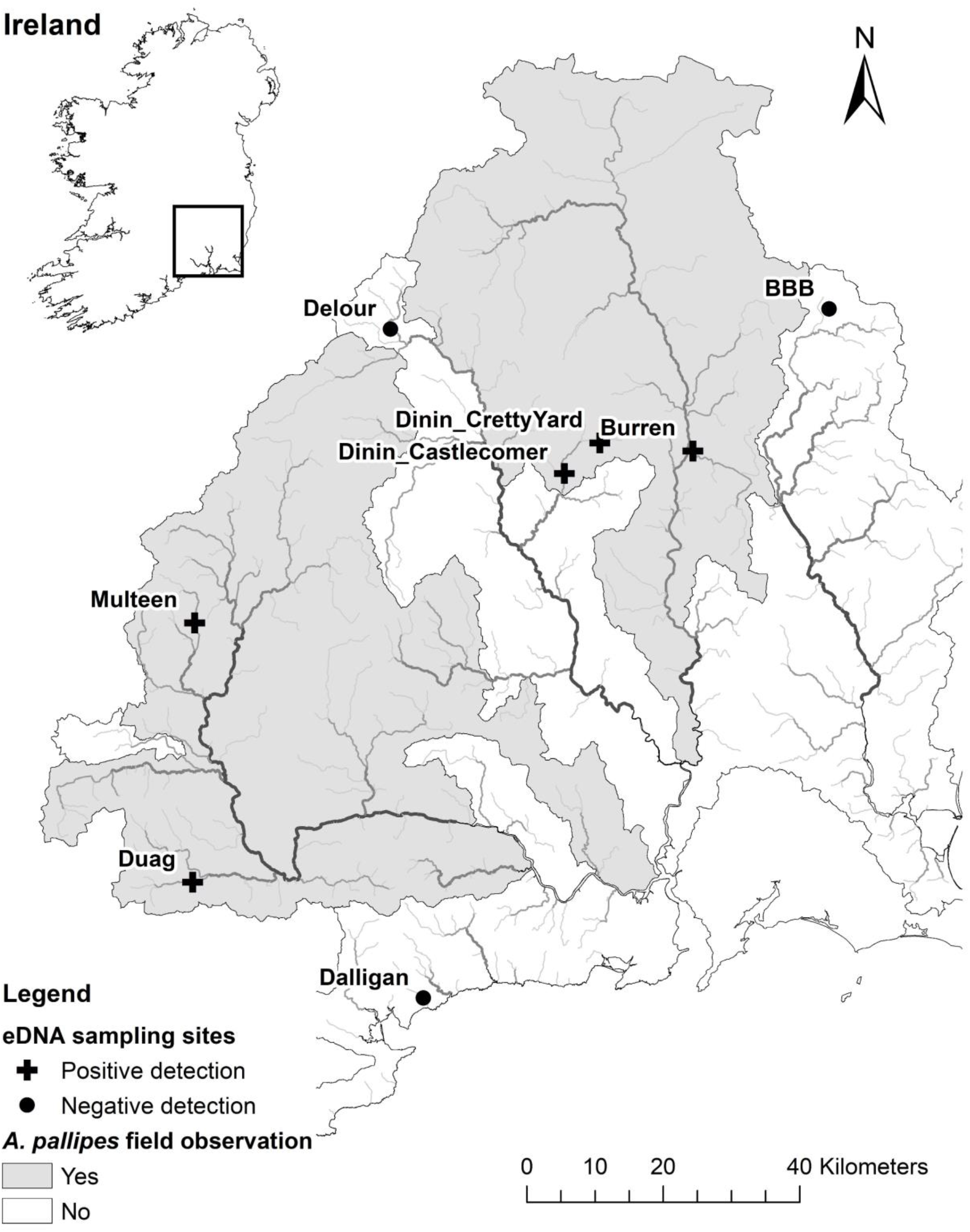
Map showing the sampling locations for the eDNA analysis. Crosses indicate sites where a positive detection of *A. pallipes* was made in at least one field replicate. Circles indicate sites where no positive detection of *A. pallipes* was made in all field replicates. River subcatchments in which field observations of *A. pallipes* were made between the years 2007 and 2016 are highlighted in grey.

When compared with the EPA’s *A. pallipes* observation data, the eDNA results reflected what was observed in the field. In all subcatchments where *A. pallipes* was observed by EPA staff, the eDNA results revealed a positive detection, and in all subcatchments where *A. pallipes* was not observed by EPA staff, the eDNA results revealed a negative detection (Fig. 1). Furthermore, it is worth noting that the eDNA results also reflected the expected distribution of *A. pallipes* based on environmental conditions. In all sites where *A. pallipes* was not detected by either field observation or eDNA (Delour, Brown’s Beck Brook and the Dalligan), the dominant bedrock was siliceous. *A. pallipes* are known to occur in waterbodies with underlying calcareous bedrock rather than siliceous bedrock (Lucey and McGarrigle 1987).

It was not possible in this study to ascertain the impact of the river obstacles on *A. pallipes* using eDNA. This was because in most sites, *A. pallipes* were either detected both above and below the obstacle, or as observed at the Dinin (Castlecomer) site, above the obstacle only. However, studies where crayfish have been physically marked or tagged have shown that river obstacles can hinder their upstream migration (Kerby et al. 2005, Bubb et al. 2008, Rosewarne et al. 2013). Environmental DNA assays for diadromous species such as sea lamprey *P. marinus* (Gustavson et al. 2015) or Chinook salmon *Oncorhynchus tshawytscha* (Laramie et al. 2015) are likely to be better indicators for assessing the migratory impact of river obstacles. Alternatively, eDNA could be used for monitoring whether invasive species have moved above the barriers designed to block them (Cowart et al. 2018). Nonetheless, the present research shows that eDNA is a reliable tool for detecting *A. pallipes* in river systems. Furthermore, while the focus of this study was in lotic systems, the assay could be readily deployed in lentic systems such as lakes or reservoirs where field sampling may be more challenging.

To conclude, this assay provides an alternative, rapid method for locating *A. pallipes* in both lentic and lotic ecosystems, allowing for more targeted, efficient surveying strategies. Considering the current crayfish plague outbreak in Ireland (National Biodiversity Data Centre 2018), and indeed other countries where it is present, there is an urgent need for a non-invasive sampling methodology that will minimise further spread of the pathogen. Large scale deployment of this assay would enable managers to rapidly establish, with confidence, the distribution of *A. pallipes*, which would support the designation of new conservation areas for the species.

## Acknowledgements

This research was funded by the Irish Environmental Protection Agency (Reconnect project - 2015-W-LS-8). The authors would also like to thank the Irish Environmental Protection Agency for providing the data on *A. pallipes* distributions.

